# Three-Dimensional Quantitative Structure-Activity Relationships and Molecular Dynamic Simulations Studies to Discover Aurora Kinase-B Inhibitors

**DOI:** 10.1101/2024.07.29.605534

**Authors:** Selvam Athavan Alias Anand, Senthamaraikannan Kabilan

## Abstract

The serine-threonine kinase gene Aurora–B kinase plays a critical role in spindle assembly, chromosome alignment, mitotic checkpoint activation, and cytokinesis. The overexpression of Aurora-B causes errors in cell division and multinucleation in centrosome numbers leads to cancer. Three–dimensional Quantitative Structure-Activity Relationship studies were conducted on known inhibitors to find valid pharmacophore hypotheses. The five features hypothesis AADRR with better parameters using partial least square analysis has been selected for virtual screening. Molecular docking was applied to find the binding mode interactions of ligands with the Aurora-B binding pocket. Lys 106, Ala 157, Glu 161, and Phe 219 were identified as crucial residues that formed several interactions with the ligands which are essential for Aurora–B inhibition. After the different levels of screening, five compounds from the National Cancer Institute database were acknowledged as novel inhibitors of Aurora–B. The active site interactions of the protein-ligand complex were examined by molecular dynamics simulation studies.

**Graphical Abstract:** 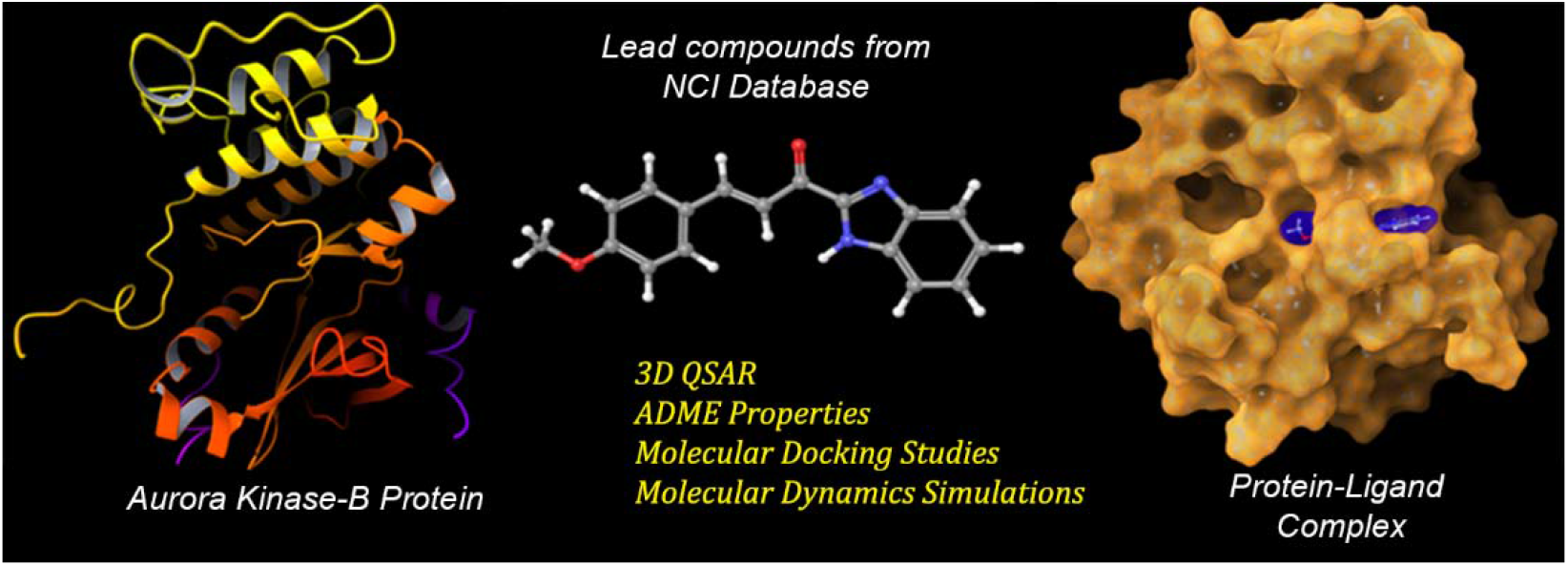

## 1. Introduction

The Aurora proteins family of serine/threonine kinases plays key roles in the event of mitosis, chromosome segregation, and cytokines [1]. During mitosis, the change in the organization of DNA due to incorrect segregation of chromosomes and unbalanced control in cell cycle checkpoints are the major characteristics of most tumour cells. The Aurora kinases are highly expressed in mitotically active cells such as the spleen, thymus, testis, bone marrow, intestine, and fetal liver [2]. Homo sapiens express three Aurora kinase paralogues Aurora–A,–B, and –C [3,4,5]. All three kinases show 67–76 % of sequence identity [6,7]. Aurora–A primarily localizes to spindle poles/centrosomes, and its disruption leads to centrosome separation and mitotic errors [8,9]. In contrast, Aurora–B, known as a chromosomal passenger protein, initially resides at chromosome centromeres during early mitosis, shifting to microtubules during anaphase [10,11]. Aurora–B plays a crucial role in chromosome segregation, alignment, spindle-checkpoint function, and cytokinesis, interacting with partners like INCENP, survivin, and borealin [12]. Aurora–C’s precise function in mammals remains unclear, but it complements Aurora–B and aids cytokinesis [13]. Dysregulation of Aurora–A and –B is associated with various cancers, making Aurora–B a promising oncogene target [14-19]. Notable Aurora kinase inhibitors include Hesperadin [20], ZM447439 [21], and VX680 [22], with Hesperadin and VX680 inhibiting all three kinases. ZM447439 selectively targets proliferating tumour cells during mitosis, sparing non-proliferating cells. VX680 exhibits promising results in arresting tumour cell proliferation in vivo and in animal models.

In the quest to discover novel scaffold inhibitors for a range of targets, various in-silico approaches, including 3D–QSAR, play a pivotal role [23,24]. In the case of inhibiting Aurora–B activity, a ligand-based pharmacophore model was employed to pinpoint the essential chemical features shared by effective inhibitors. To construct these hypotheses, a collection of known Aurora–B inhibitors was sourced from the Binding Database website (www.bindingdb.org/). The selection of the most robust hypothesis relied on rigorous statistical criteria, including the coefficient of determination (R^2^), cross-validated correlation coefficient (Q^2^), standard deviation (SD), variance ratio (F), and root mean square error (RMSE) values. This selected hypothesis then served as a 3D query to initiate the initial filtration process in virtual screening. Subsequently, the screened compounds underwent further refinement to ensure optimal pharmacokinetic properties, after which they were subjected to molecular docking studies to identify the best-fit molecules. The ultimate decision on lead compounds was made through a comprehensive comparison of docking analysis and pharmacokinetic properties. These chosen leads were then subjected to rigorous evaluation through molecular dynamics (MD) simulations to elucidate their binding mode interactions. This study’s advanced in-silico methods can accelerate drug discovery, particularly for Aurora–B inhibitors, through rigorous statistical analysis, virtual screening, and molecular dynamics simulations, providing valuable insights into lead compound identification and evaluation.

## 2. Materials and Methods

### 2.1 Protein Preparation

Ligand-based pharmacophore modelling studies were carried out by using the PHASE application [25] implemented with Maestro [26]. The 3D coordinates of the crystallographic structure of the human Aurora–B: INCENP (PDB ID: 4AF3) complex [27] were downloaded from Brookhaven Protein Data Bank (www.rscb.com). The protein complex was pre-processed and prepared by Protein Preparation Wizard [28] in Maestro of Schrödinger software. The minimization of the complex was continued using the OPLS–2005 (Optimized Potential for Liquid Simulations) force field [29] until the root mean square deviation (RMSD) reached the value of 0.3 Å. The protein underwent preprocessing, involving the correction of heavy atoms, water molecules, cofactors, and metal ions. Additionally, missing hydrogen atoms, side chains, and protons were added. The Schrodinger suite’s EPIK and IMPREF programs were employed for further refinement and minimization. To define the active site vicinity, the receptor grid generation module was utilized, creating a grid that encapsulated the ligand’s surroundings. The resulting workspace configuration, appearing as a centroid in a cubic shape, precisely delineated the protein’s active site. The molecular docking studies of hit molecules were performed by GLIDE [30]. MD simulation studies were carried out by the DESMOND [31] module.

### 2.2. Pharmacophore Modelling

A total of 58 known inhibitors of Aurora–B with quantitative bioactivity data were used to design the pharmacophore model. The inhibitors were randomly divided into the training set and test set for structure-based pharmacophore generation and validation. Highly active and least active compounds were included in both training and test sets. The critical information on pharmacophore design was provided by training set ligands.

The five steps for pharmacophore modelling are given below:

#### 2.2.1 Ligand Preparation

The 2D structures of compounds were imported from the project table to the Develop Common Pharmacophore Hypotheses panel. The structures were minimized and geometrically refined using the LIGPREP [32, 33] module which neutralizes the ionized structures to a pH7 (neutral) and generates possible stereoisomers. Conformers were generated using the rapid torsional angle search method ‘ConfGen’ with distance-dependent dielectric solvation treatment and OPLS–2005 force field incorporated in PHASE. The molecular docking simulations using an implicit solvent model with a distance-dependent dielectric (GB/SA) approach. The simulations employed a cutoff of 1 Å root mean square deviation for interactions and consisted of 1000 iterations, with water as the implicit solvent. For each structure, a maximum of 1000 conformers were generated using 100 steps of pre-process minimizations and 50 steps of post-process minimizations. The maximum energy difference for a set of conformers of each molecule is 10 kcal/mol. The active and inactive ligands were assigned by giving appropriate values in the activity threshold.

#### 2.2.2 Creating Pharmacophore Sites

This step is used to set pharmacophore features to create sites for all ligands. Six pharmacophore features provided by PHASE are hydrogen bond acceptor (A), hydrogen bond donor (D), hydrophobic group (H), negatively ionisable (N), positively ionisable (P), and aromatic ring (R). All six features were utilized for the creation of pharmacophore sites.

#### 2.2.3 Finding Common Pharmacophore

Common pharmacophores were identified from the set of variants using a tree-based partition technique with a maximum depth of 5 and a minimum intersite distance is 2.0 Å. The initial and final box sizes are 32.0 Å and 1.0 Å respectively, which all activities should match. The Common Pharmacophore Hypotheses (CPHs) were generated by varying the maximum and minimum number of sites and the number of matching active groups.

#### 2.2.4 Scoring Hypotheses

The CPHs were examined using the scoring procedure to find the best alignment of active set molecules. The scoring process provides hypotheses with different features based on ranking. The scores displayed in the hypotheses table are used to choose the appropriate hypothesis for further investigation.

#### 2.2.5 Building QSAR Model

In QSAR modelling, the data set was divided into a training set (70%) and a test set (30%) for the selected hypothesis. PHASE provides atom-based QSAR model and a pharmacophore features-based QSAR model for the structural components that form the basis of the model. We used an atom-based QSAR model, in which the structural components of ligands were represented by van der Waals models of atoms in the ligands. The atoms occupied in the same region of the state are distinguished by six classes:

- D – Hydrogen–bond donor (hydrogen bonded to N, O, P, S)
- H – Hydrophobic or nonpolar (C, H–C, halogens)
- N – Negative ionic (formal negative charge)
- P – Positive ionic (formal positive charge)
- W – Electron withdrawing (N, O includes hydrogen–bond acceptor)
- X – Miscellaneous (all other groups)

The partial least square (PLS) regression was carried out for the QSAR model using PHASE with a maximum of N/5 factors (where, N = number of ligands in the training set). The accuracy of the models increased with the increasing number of PLS factors until overfitting started to occur [34,35]. Three PLS factors were generated for all the hypotheses with a grid spacing of 1 Å and the best model was selected on the basis of significant statistical data such as R^2^, Q^2^, SD, RMSE, F, Pearson R, and stability values for virtual screening.

#### 2.3 Virtual Screening

The validated hypothesis is used as a query to search the novel molecules such as Aurora–B inhibitors. The NCI and Maybridge databases were explored for desired chemical structures. In accordance with Lipinski’s rule of five [36,37], the hit molecules obtained were then screened by using the QIKPROP module [38] for the *in silico* determination of pharmacokinetic properties such as absorption, distribution, metabolism, and excretion (ADME) to make them more drug-like. The molecules with drug-likeness were subjected to molecular docking to find the best-fit molecules in the active site of the Aurora–B protein. The “Glide” software offers three distinct levels of docking methodologies: High Throughput Virtual Screening (HTVS), Standard Precision (SP), and Extra Precision (XP). Within Schrodinger’s Glide module, HTVS docking was employed to predict protein-ligand binding modes and rank ligands utilizing empirical scoring functions. The subsequent SP docking step with Glide refines the rankings of ligands initially docked in HTVS mode. Moreover, the XP Glide approach, following SP docking, conducts a meticulous refinement of docking modes through an anchor and grow algorithm. Initial filtering of molecules was done by using high–throughput virtual screening (HTVS); next the standard precision (SP), and finally, the top hit molecules were selected for the extra precision (XP) mode of docking [39]. The lead molecules were selected based on the comparison of Glide–score and pharmacokinetic properties.

#### 2.4 Molecular Dynamics (MD) Simulations

The molecules with top-ranked G–score and acceptable parameters of pharmacokinetic properties were selected for MD simulations study using DESMOND with OPLS 2005 force field. The water molecules were placed around the protein-ligand complex with a predefined TIP3P water model [40] in an orthorhombic box. The overall charge was neutralized by adding salt counter–ions. The temperature and pressure were kept constant at 300 K and 1.01325 bar using the Nose-Hoover thermostat [41] and Martyna–Tobias–Klein barostat [42] methods. The simulations were performed using an NPT ensemble by considering the number of atoms, pressure, and timescale. During simulations, the long-range electrostatic interactions were calculated using the Particle-Mesh-Ewald method [43,44].

## 3. Results and Discussion

### 3.1 QSAR Pharmacophore modelling

A dataset of 40 ligands was selected for the training set and 18 ligands for the test set randomly. The chemical structure of training and test set ligands were given in the supplementary data (Table S1 and Table S2). The ligands with -logIC_50_ higher than 7.7 were considered ‘active’ and lower than 6.5 were considered ‘inactive’, whereas the ligands with in-between -logIC_50_ values were considered as moderately active for the creation of CPHs. After ligand preparation, the scoring hypothesis was done by keeping the RMSD value below 1.2 Å and the vector score above 0.5. With the use of the tree-based partition technique, the pharmacophore identification results in 24 different variant hypotheses. The best hypothesis was selected based on the alignment of site points and vectors, volume overlap, selectivity, number of ligands matched, relative conformational energy, and activity. The hypothesis with five features AADRR was considered the best model based on R^2^, SD, F, Q^2^, RMSE, stability, and Pearson R values. The hypothesis includes two hydrogen bond acceptors (A), one hydrogen bond donor (D), and two aromatic rings (R) a total of five pharmacophoric features. The actual and predicted IC_50_ values of training and test set molecules with their fitness are presented in Table 1 and the plot of actual versus predicted IC_50_ for training and test set ligands is shown in Fig. 1.

**Table 1.** The fitness and activity data of training and test set molecules. The actual and predicted IC_50_ values of training and test set molecules with their fitness are presented in this table.

| Ligand No. | QSAR Set | $-\log\text{IC}_{50}$ | Predicted Activity for PLS Factor 3 | Residuals | Fitness |
| --- | --- | --- | --- | --- | --- |
| 1 | Training | 9.398 | 9.31 | -0.088 | 0.99 |
| 2 | Training | 9.301 | 9.21 | -0.091 | 1.04 |
| 3 | Test | 9.222 | 9.19 | -0.032 | 1.02 |
| 4 | Training | 9.155 | 9.19 | 0.035 | 1.05 |
| 5 | Test | 9.097 | 9.18 | 0.083 | 1.06 |
| 6 | Training | 9.000 | 9.17 | 0.170 | 1.05 |
| 7 | Training | 8.022 | 8.01 | -0.012 | 2.68 |
| 8 | Training | 7.971 | 7.82 | -0.151 | 1.57 |
| 9 | Test | 7.959 | 7.62 | -0.339 | 2.66 |
| 10 | Test | 7.886 | 7.89 | 0.004 | 1.64 |
| 11 | Test | 7.854 | 7.36 | -0.494 | 2.12 |
| 12 | Test | 7.810 | 7.35 | -0.460 | 1.21 |
| 13 | Test | 7.770 | 7.62 | -0.150 | 0.93 |
| 14 | Training | 7.745 | 7.69 | -0.055 | 2.2 |
| 15 | Training | 7.721 | 7.78 | 0.059 | 1.29 |
| 16 | Test | 7.703 | 7.70 | -0.003 | 3 |
| 17 | Training | 7.699 | 7.69 | -0.009 | 2.61 |
| 18 | Training | 7.678 | 7.61 | -0.068 | 1.68 |
| 19 | Training | 7.635 | 7.66 | 0.025 | 1.28 |
| 20 | Training | 7.602 | 7.64 | 0.038 | 2.65 |
| 21 | Training | 7.600 | 7.65 | 0.050 | 1.76 |
| 22 | Training | 7.592 | 7.64 | 0.048 | 1.44 |
| 23 | Training | 7.523 | 7.53 | 0.007 | 1.19 |
| 24 | Training | 7.394 | 7.36 | -0.034 | 1.61 |
| 25 | Training | 7.364 | 7.33 | -0.034 | 1.71 |
| 26 | Training | 7.299 | 7.35 | 0.051 | 1.51 |
| 27 | Test | 7.300 | 7.17 | -0.130 | 1.45 |
| 28 | Training | 7.276 | 7.19 | -0.086 | 1.03 |
| 29 | Test | 7.247 | 7.20 | -0.047 | 1.64 |
| 30 | Training | 7.102 | 7.10 | -0.002 | 1.12 |
| 31 | Training | 7.077 | 7.12 | 0.043 | 1.46 |
| 32 | Training | 7.036 | 7.07 | 0.034 | 1.52 |
| 33 | Training | 7.027 | 7.07 | 0.043 | 1.48 |
| 34 | Test | 7.004 | 6.54 | -0.464 | 1.22 |
| 35 | Training | 6.975 | 6.99 | 0.015 | 1.05 |
| 36 | Test | 6.910 | 6.97 | 0.060 | 1.02 |
| 37 | Training | 6.857 | 6.71 | -0.147 | 1.26 |
| 38 | Training | 6.842 | 6.85 | 0.008 | 1.61 |
| 39 | Test | 6.800 | 6.90 | 0.100 | 1.34 |
| 40 | Training | 6.796 | 6.81 | 0.014 | 1 |
| 41 | Test | 6.759 | 7.13 | 0.371 | 1.62 |
| 42 | Training | 6.745 | 6.88 | 0.135 | 1.54 |
| 43 | Training | 6.719 | 6.67 | -0.049 | 2.19 |
| 44 | Training | 6.712 | 6.67 | -0.042 | 1.81 |
| 45 | Training | 6.699 | 6.78 | 0.081 | 1.18 |
| 46 | Training | 6.678 | 6.64 | -0.038 | 1.2 |
| 47 | Test | 6.646 | 6.97 | 0.324 | 1.4 |
| 48 | Training | 6.636 | 6.64 | 0.004 | 1.25 |
| 49 | Test | 6.635 | 6.93 | 0.295 | 1.39 |
| 50 | Test | 6.600 | 7.03 | 0.43 | 1.22 |
| 51 | Training | 6.400 | 6.42 | 0.02 | 0.49 |
| 52 | Training | 6.312 | 6.16 | -0.152 | 1.76 |
| 53 | Test | 6.232 | 6.14 | -0.092 | 1.76 |
| 54 | Training | 6.111 | 6.18 | 0.069 | 1.78 |
| 55 | Training | 6.107 | 6.18 | 0.073 | 1.74 |
| 56 | Training | 6.013 | 6.12 | 0.107 | 1.71 |
| 57 | Training | 6.000 | 5.96 | -0.04 | 1.08 |
| 58 | Training | 5.900 | 5.91 | 0.01 | 0.95 |

**Fig. 1.**
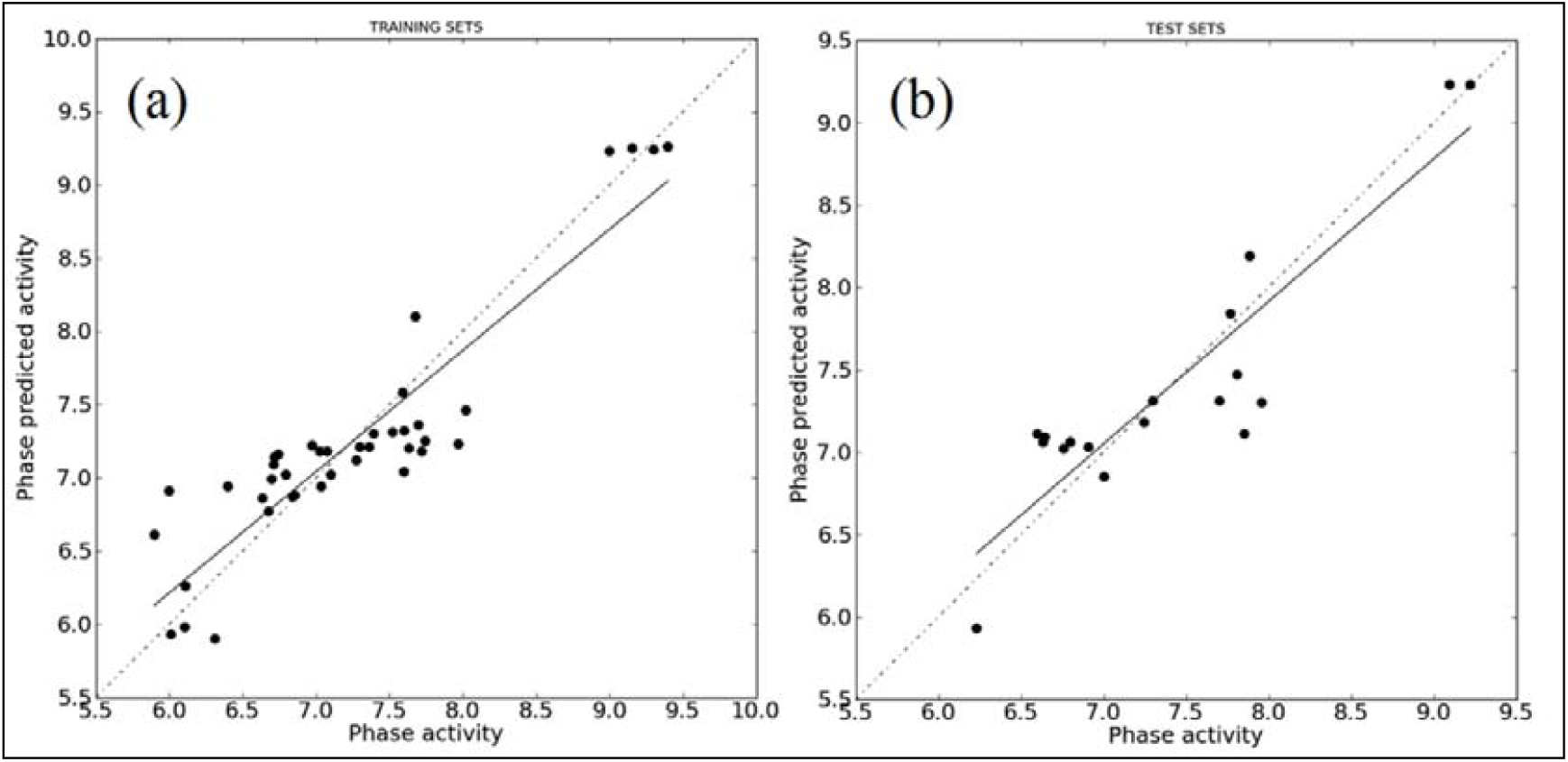
The plot of actual versus predicted IC_50_ for (a) training set and (b) test set ligands

According to Tropsha [45], high r^2^ is a necessary but not sufficient condition for a QSAR model. It is also supported by RMSE and Pearson R values. Training set compounds were aligned in the AADRR pharmacophore model and analyzed with three PLS factors in PHASE. The QSAR results of AADRR are R^2^=0.971, Q^2^=0.907, Fit value=403, and SD value =0.154, indicating that this model is a good predictive hypothesis. Therefore AADRR model was selected for QSAR analysis. The statistical parameters of AADRR are given in Table 2. The correlation squared value for the test set molecules (R^2^=0.81) confirms the better predictability of the QSAR model for the test set molecules. The spatial arrangement of the five featured pharmacophore models along with their distance is shown in Fig. 2.

**Table 2.** Statistical parameters of the Best Hypothesis. The QSAR results of the AADRR model showed R^2^=0.971, Q^2^=0.907, Fit value=403, and SD value =0.154, Pearson-R = 0.952, indicating that this is a good predictive hypothesis.

| Hypothesis ID | PLS Factors | SD | $R^2$ | F | P | Stability | RMSE | $Q^2$ | Pearson-R |
| --- | --- | --- | --- | --- | --- | --- | --- | --- | --- |
| AADRR.94 | 1 | 0.3668 | 0.8284 | 183.5 | 3.999 e-16 | 0.9384 | 0.357 | 0.804 | 0.8989 |
|  | 2 | 0.2337 | 0.9322 | 254.3 | 2.389 e-22 | 0.8595 | 0.2609 | 0.8953 | 0.9463 |
|  | 3 | 0.1547 | 0.9711 | 403 | 9.626 e-28 | 0.7662 | 0.245 | 0.9077 | 0.9529 |

SD – Standard deviation of the regression, $R^2$ – Coefficient of determination, F – variance ratio, P – Significance level of the variance ratio, RMSE – Root mean square error value, $Q^2$ – Cross-validated correlation coefficient, Pearson R – correlation between predicted and observed activity for test set.

**Fig. 2.**
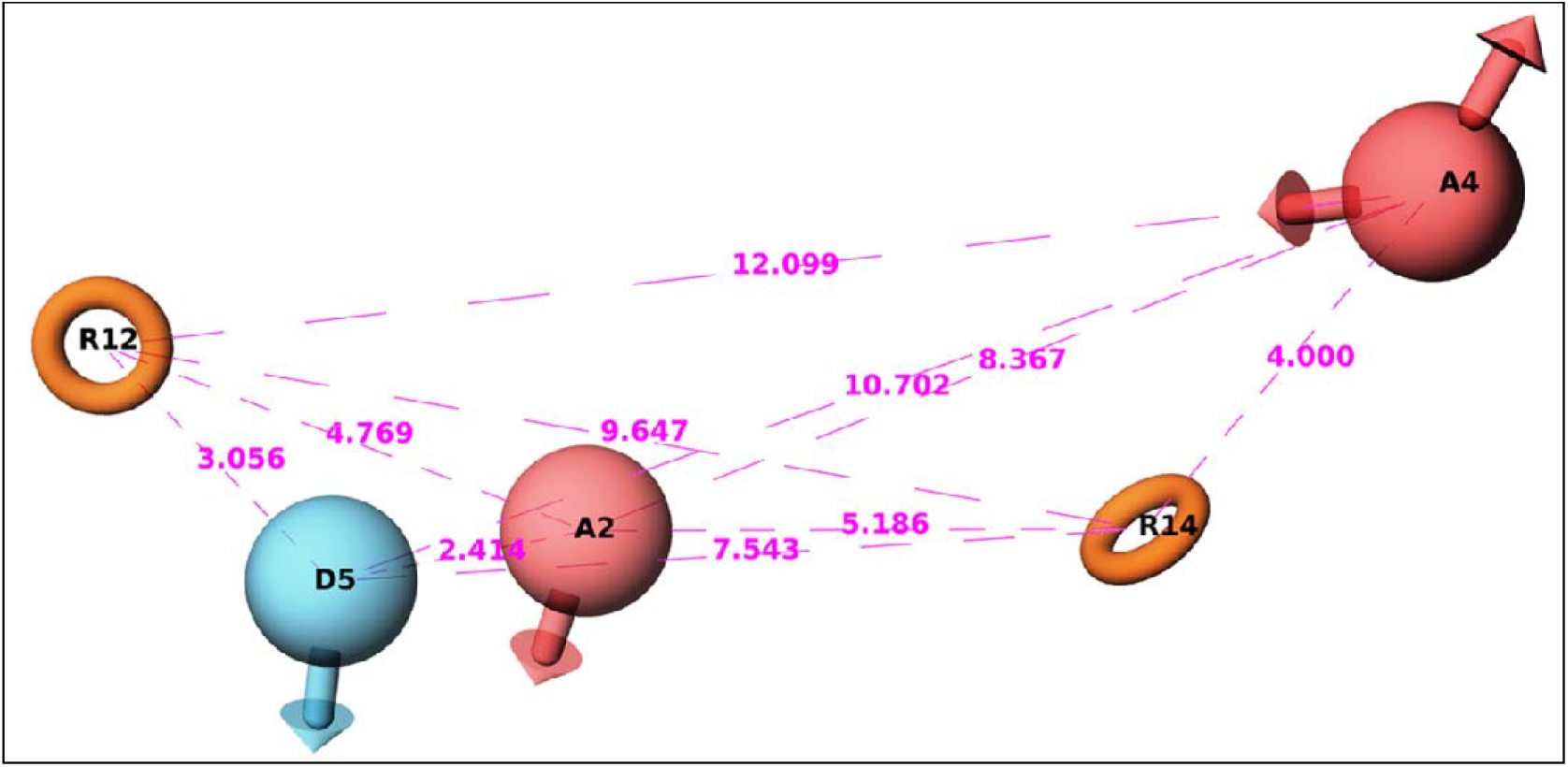
The spatial arrangement of AADRR model with two hydrogen–bond acceptors, one hydrogen–bond donor and two aromatic rings.

### 3.2 Virtual Screening

In this study, a total of 320,078 hit molecules were retrieved from two databases: the NCI database (265,242 compounds) and the Maybridge database (54,836 compounds). The initial screening process utilized ‘Pharmacophore Matching’ using the AADRR model to select the top 1,000 compounds. Subsequently, these 1,000 compounds underwent further scrutiny, applying Lipinski’s rule of five using the QIKPROP program, resulting in the identification of 822 promising compounds. These 822 compounds with good and moderate parameters of pharmacokinetic (ADME) properties then underwent a comprehensive docking analysis employing three different approaches: High Throughput Virtual Screening (HTVS), Standard Precision (SP), and Extra Precision (XP). Molecular Docking The important factor in finding potent inhibitors is to know about the residues present in the active site. Initially, Aurora–B inhibitors: Hesperadin, ZM447439, and VX–680 were docked with human Aurora–B: INCENP protein to know the critical amino acids that play a major role in the active site. The results revealed that Ala157 and Lys106 are important residues for successful binding. Lipinski’s rule filtered molecules (N=822) were subjected to rapid screening by high–throughput virtual screening (HTVS). In agreement with the docking score ranking, the top molecules were selected for standard precision (SP) docking. Further, the first 58 compounds were selected after extra precision (XP) docking. The flow chart of the virtual screening technique used is presented in Fig. 3. Finally, by the evaluation of G– score, pharmacokinetic properties, and the interactions with critical amino acids, five compounds with NCI ID 695163 (compound **1**), 327359 (compound **2**), 721045 (compound **3**), 711797 (compound **4**), 104546 (compound **5**), were identified as lead compounds (Fig. 4) with different scaffolds from NCI database. The interactions between Aurora–B protein and lead compounds are shown in Fig. 5. The pharmacokinetic properties of lead compounds are given in Table 3. The lead compounds showed no violation of Lipinski’s rule of five and the results of other pharmacokinetic properties were also satisfactory.

**Table 3.** The ADME properties of the lead compounds.

| S.No. | Drug-like Parameters | Pharmacokinetic Properties of the Lead Compounds with NCI ID |  |  |  |  |
| --- | --- | --- | --- | --- | --- | --- |
|  |  | 695163 | 327359 | 721045 | 711797 | 104546 |
| 1 | mol_MW<sup>a</sup> | 278.31 | 330.419 | 447.831 | 335.318 | 330.345 |
| 2 | Donor HB<sup>b</sup> | 1 | 2 | 1 | 1 | 3 |
| 3 | Accpt HB<sup>c</sup> | 4.25 | 4.95 | 8.45 | 8 | 5.5 |
| 4 | QLogPo/w<sup>d</sup> | 3.148 | 3.132 | 3.005 | 1.554 | 2.338 |
| 5 | **Rule of Five** | 0 | 0 | 0 | 0 | 0 |
| 6 | Percent Human Oral Absorption<sup>e</sup> | 100 | 96.956 | 89.088 | 84.471 | 84.442 |
| 7 | QLogS<sup>f</sup> | -4.055 | -4.125 | -4.928 | -2.975 | -3.97 |
| 8 | QLogHERG<sup>g</sup> | -5.971 | -5.916 | -6.663 | -5.473 | -6.083 |
| 9 | QLogBB<sup>h</sup> | -0.56 | -0.909 | -1.26 | -1.012 | -1.298 |

<sup>a</sup> Molecular weight of the compound (< 500); <sup>b</sup> Number of hydrogen bond donor (< 5); <sup>c</sup> Number of hydrogen bond acceptor (< 10); <sup>d</sup> Partition coefficient value between octanol and water (< 5); <sup>e</sup> Percentage of human absorption, (> 80 high, < 25 poor); <sup>f</sup> Predicted aqueous solubility; S in mol/L, (between -6.5 to 0.5); <sup>g</sup> Predicted IC<sub>50</sub> value for blockage of HERG K<sup>+</sup> channels, (below -5); <sup>h</sup> Predicted blood-brain barrier permeability, (between -3 to 1.2)

**Fig. 3.**
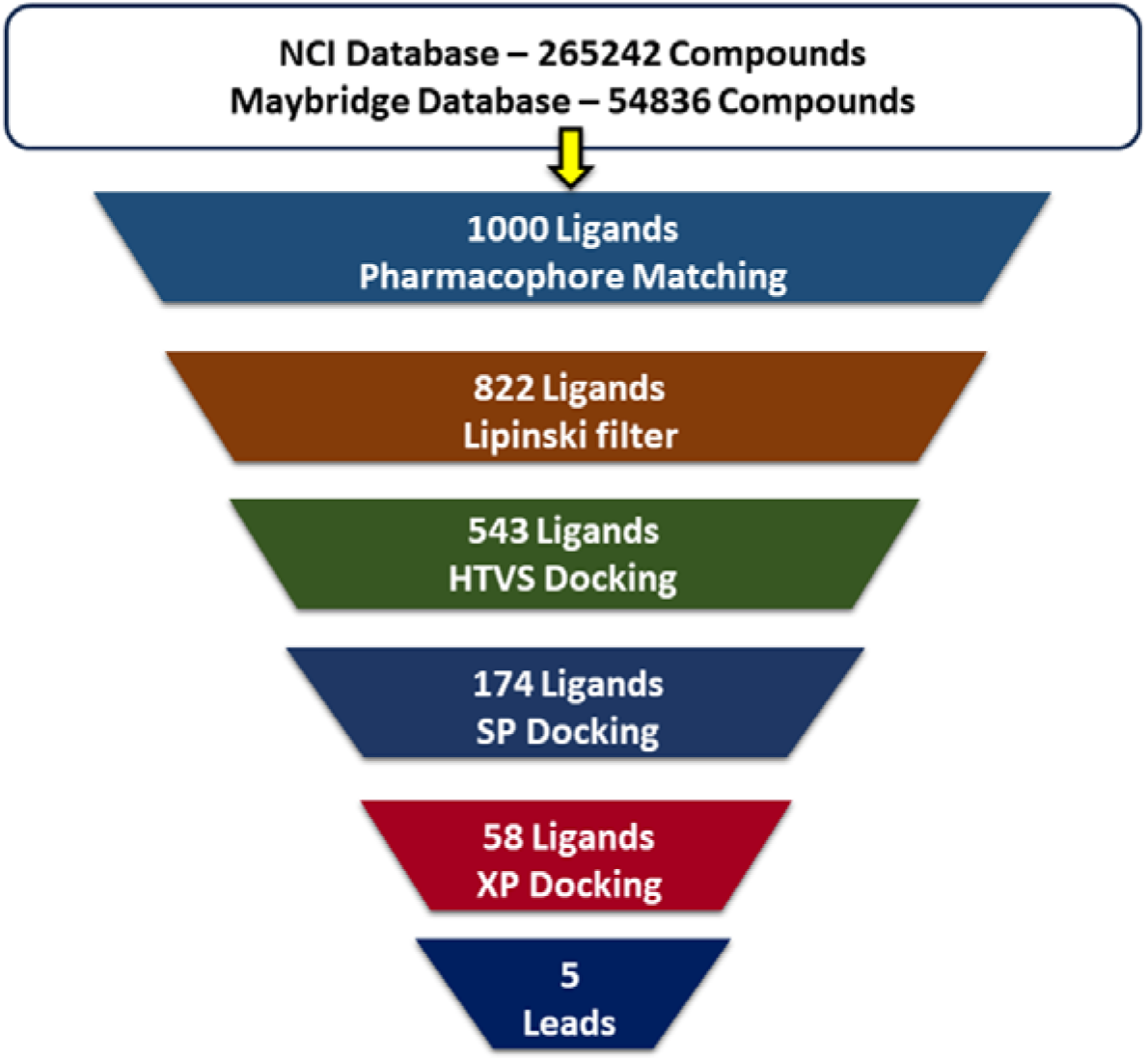
The flow chart of virtual screening technique. After the screening methods five lead compounds were identified from the binding database and used for the analysis.

**Fig. 4.**
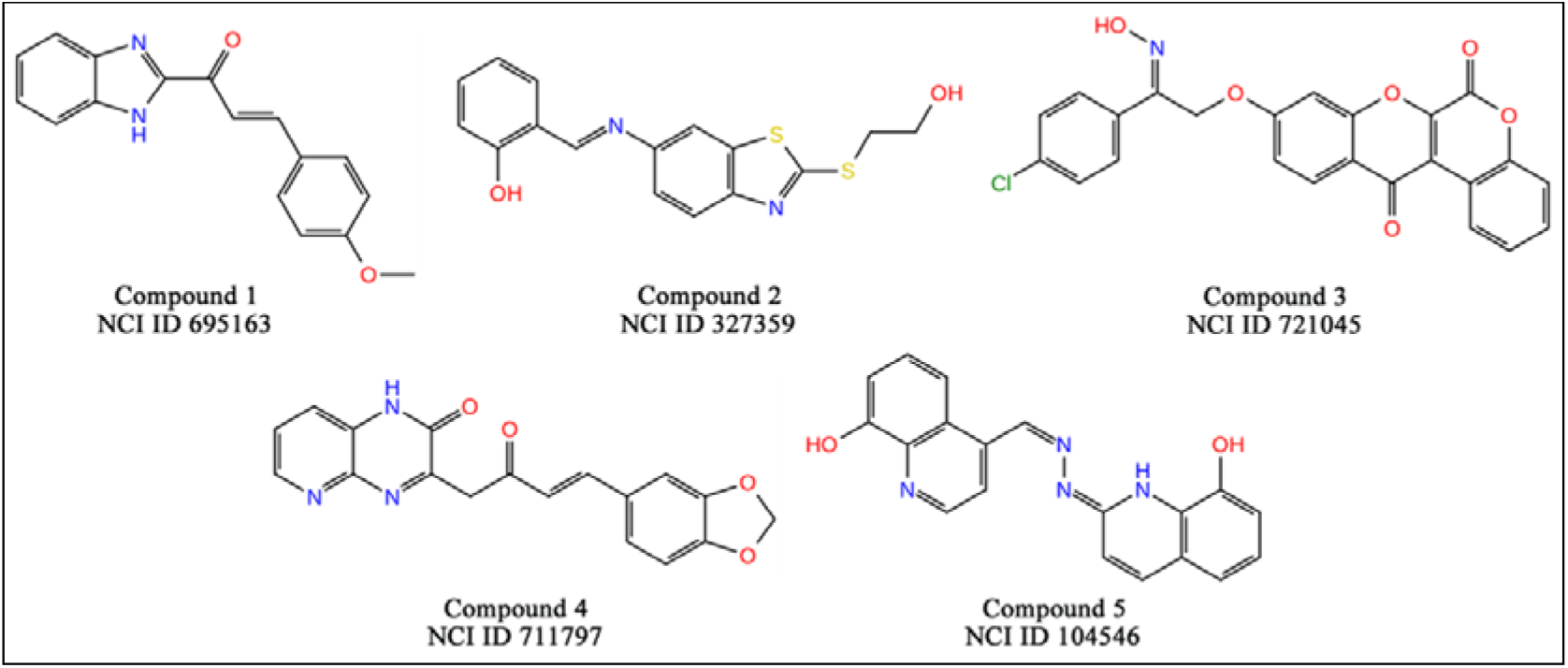
The chemical structures of lead compounds

**Fig. 5.**
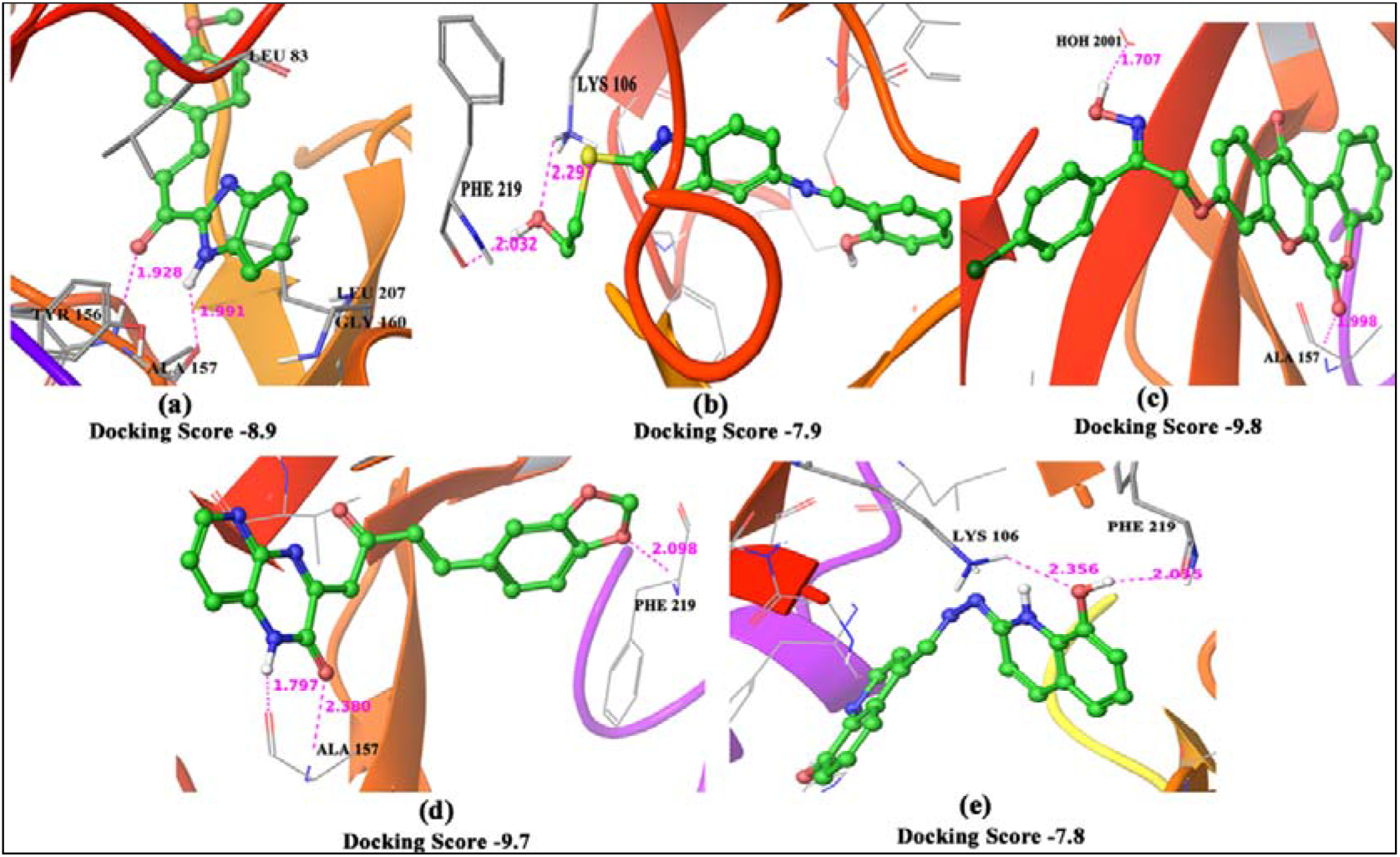
Docking poses of Aurora–B protein (PDB ID: 4AF3) with the lead compounds. The molecular docking studies were carried out by Glide module of Schrodinger software. The protein 4AF3 downloaded from the protein data bank website (https://www.rcsb.org/) and the lead compound structures from the binding database website (http://www.bindingdb.org) (a) Compound 1, (b) Compound 2, (c) Compound 3, (d) Compound 4, (e) Compound 5

### 3.3 Molecular Dynamics Simulations

In order to clarify the interaction patterns between the Aurora–B protein and lead compounds, molecular dynamics simulations were carried out for compounds **1** to **5**. Since in MD simulations, both the protein and ligand are flexible, the adjustment of conformation gives valid information about the binding affinity of ligands. The relative RMSD was calculated to find the conformational change during simulations. The RMSD plot (Fig. 6) reveals that the structures of compound **1**, compound **4** and compound **5** were stable after 3ns simulations, whereas for compound **2** and compound **3**, the enzymes increased still 4ns to adopt optimal conformation.

**Fig. 6.**
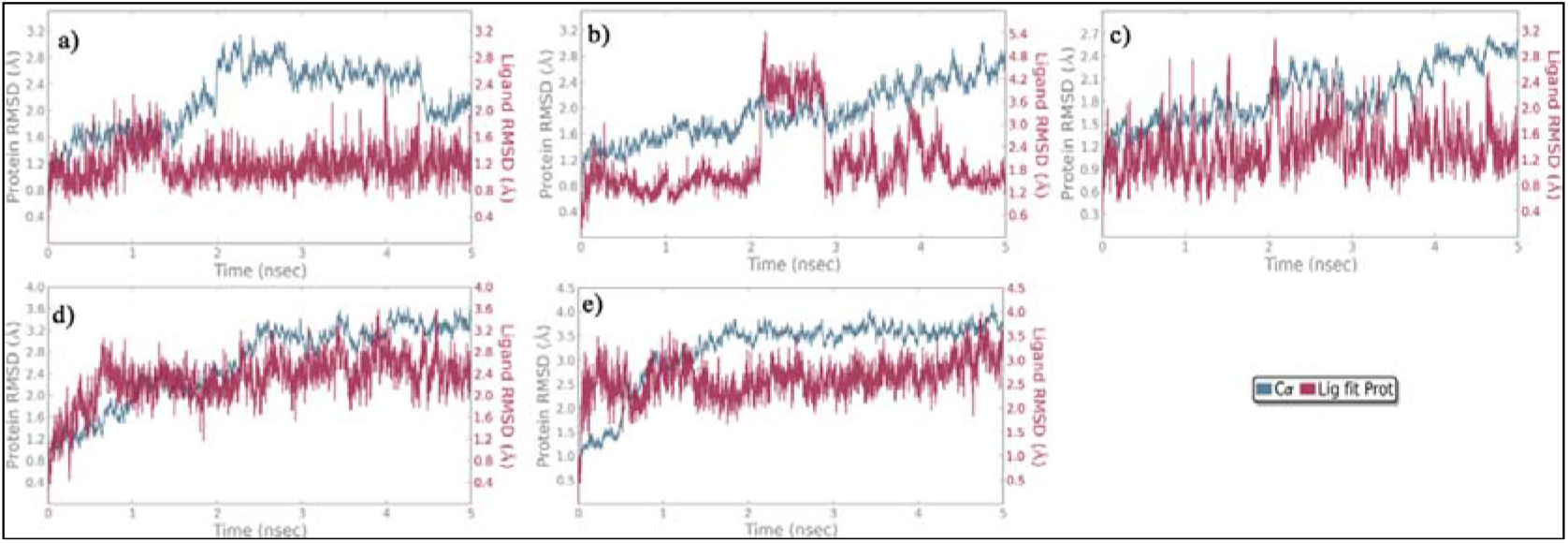
The RMSD of Aurora–B protein complex with (a) Compound 1, (b) Compound 2, (c) Compound 3, (d) Compound 4, (e) Compound 5.

It is seen that the complex of compound **5** reaches about 3.5 Å after 2 ns and the value remains unchanged throughout the simulations. This demonstrates the conformational stability of the enzyme structure. To investigate the specific binding mode, the MD simulation results were compared with docking analysis and tabulated in Table 4. The compounds **1** to **5** show strong interactions with the critical amino acids Ala157, and Lys106. The interactions of the protein with compounds **1-5** in MD simulations are shown in Fig. 7. Since there is no significant change in protein-ligand interactions, MD simulations results validate the docking analysis.

**Table 4.**
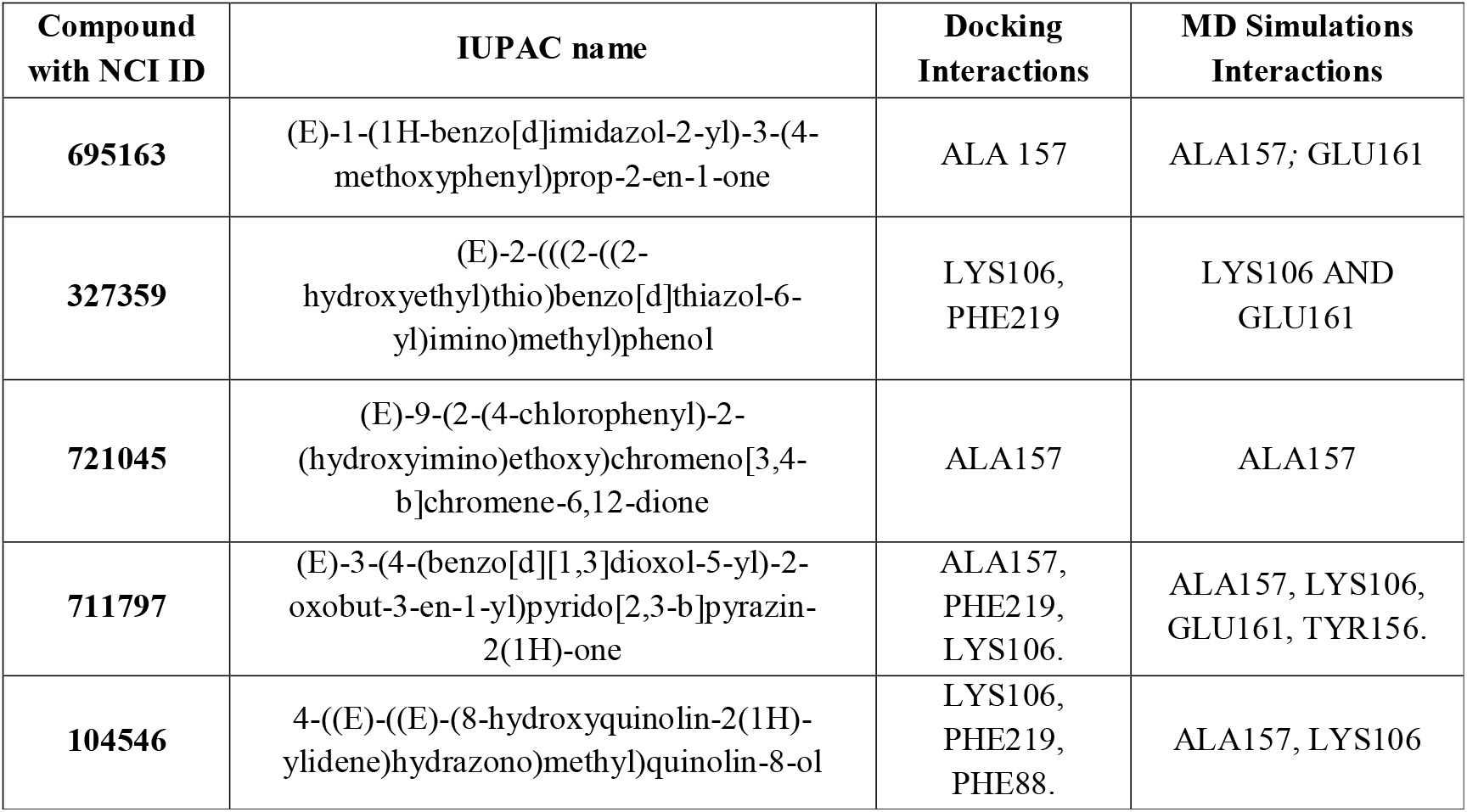
Comparison of protein-ligand interactions before and after MD simulations. The critical interactions between the five compounds and the 4AF3 protein were tested by both molecular docking and molecular dynamic simulations. The residues involve in the interactions are listed in this table.

**Fig. 7.**
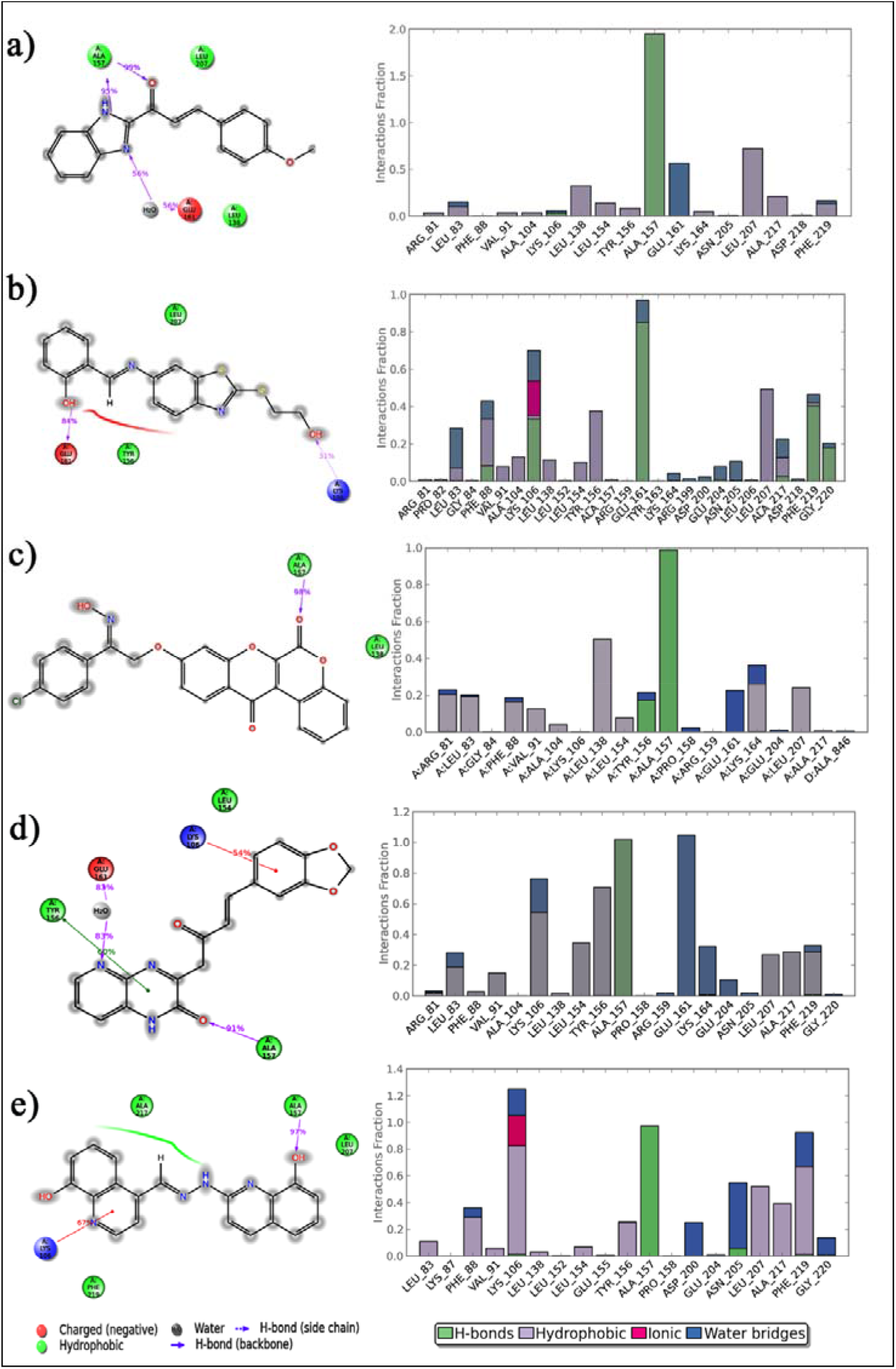
The 2D view and the histogram of protein interactions after MD with (a) Compound 1, (b) Compound 2, (c) Compound 3, (d) Compound 4, (e) Compound 5.

## 4 Conclusion

In this study, the chemical features of Aurora–B inhibitors with good statistical results have been developed by using PHASE of the Schrödinger suite. The pharmacophore hypothesis AADRR is used for the QSAR model to find potent and novel inhibitors. The crystallographic structure of the human Aurora B: INCENP complex (PDB ID: 4AF3) is used for docking and MD simulations analysis. In agreement with virtual screening, pharmacokinetic properties, and docking analysis, five compounds from the NCI database (Compound **1** – NCI ID 695163, Compound **2** – NCI ID 327359, Compound **3** – NCI ID 721045, Compound **4** – NCI ID 711797, Compound **5** – NCI ID 104546) were selected as lead compounds. To clarify the interactions between Aurora–B protein and lead compounds, MD simulations were carried out for compounds **1** to **5**. The results of docking and MD simulations studies reveal that the lead compounds have strong interactions with the critical amino acids Ala157, and Lys106 which are essential for Aurora–B inhibitory activity. Also, MD studies were used to find the conformational stability of protein complexes during simulations. From the above study, we believe that our AADRR pharmacophore model, docking, and MD simulations studies provide valid information on the structural requirements for Aurora–B kinase inhibition. In recent research [46], an investigation revealed interactions between specific residues (Leu83, Phe88, Val91, Ala157, and Glu155) within the binding pocket of the Aurora Kinase B receptor and various ligands. The researchers developed a pharmacophoric model with seven distinct features, including two hydrophobic elements, one donor, one acceptor, and three exclusion volumes. However, our study introduced a pharmacophoric model utilizing a five-feature hypothesis known as AADRR, which also exhibited better parameters when subjected to partial least square analysis. This approach led to the identification of five promising lead compounds selected from a pool of 320,000 compounds sourced from the Maybridge and NCI databases. These lead compounds were shown to engage in interactions with the Aurora-B binding pocket, underscoring their potential for further exploration. Specifically, we found that Lys 106, Ala 157, Glu 161, and Phe 219 were pivotal residues responsible for establishing multiple interactions with the ligands, further emphasizing the significance of these compounds in future research endeavours.

## Acknowledgements

We are thanking Annamalai University for providing the research facility and software support. Prof. S. Kabilan acknowledged that he received UGC BSR Faculty Fellowship.

## Declarations

### Ethical Approval

“not applicable”.

## Availability of data and materials

The data and materials are attached as supporting information.

