## Supplementary Material for "Three-Dimensional Quantitative Structure-Activity Relationships and Molecular Dynamic Simulations Studies to Discover Aurora Kinase-B Inhibitors"

**Table S1.** Chemical structures of training set compounds.

**(Source:**http://www.bindingdb.org/jsp/dbsearch/PrimarySearch_ki.jsp?tag=tg&kiunit=nM&icunit=nM&column=IC50&submit=Search&energyterm=kJ%2Fmole&target=Aurora+kinase+B)

| 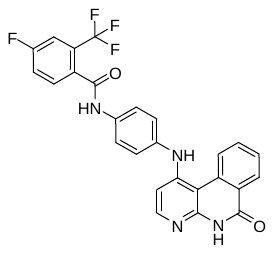  -logIC_50_ **9.398** | 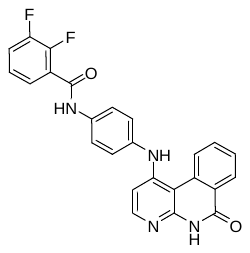 -logIC_50_ **9.301** | 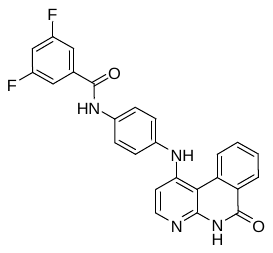  -logIC_50_ **9.155** | 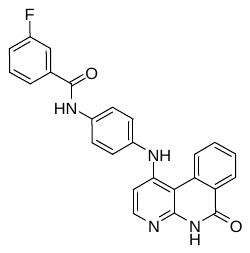 -logIC_50_ **9.000** | 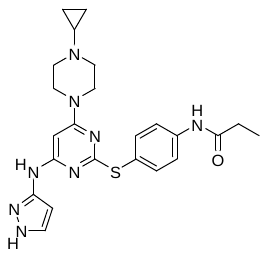 -logIC_50_ **8.022** |
| --- | --- | --- | --- | --- |
| 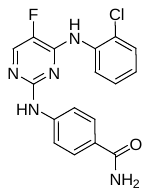  -logIC_50_**7.971** | 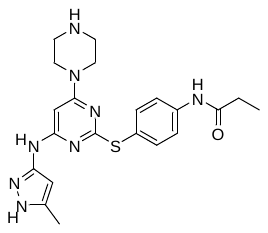 -logIC_50_ **7.745** | 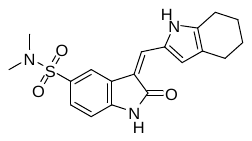  -logIC_50_ **7.721** | 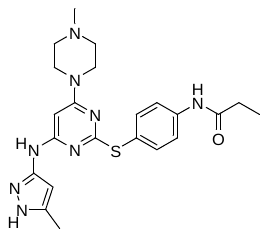 -logIC_50_ **7.699** | 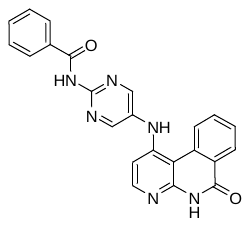 -logIC_50_**7.678** |
| 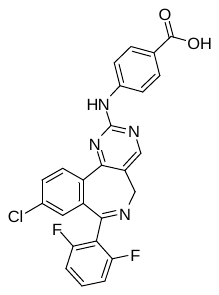  -logIC_50_ **7.635** | 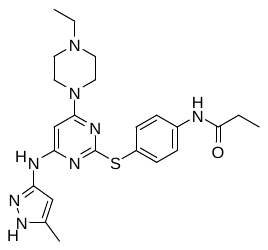 -logIC_50_ **7.602** | 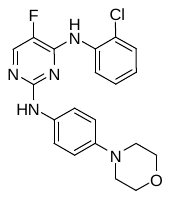  -logIC_50_ **7.600** | 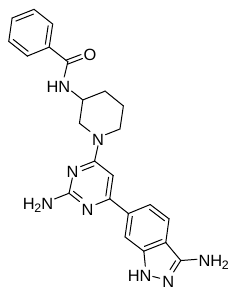  -logIC_50_ **7.592** | 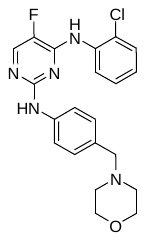  -logIC_50_ **7.523** |
| 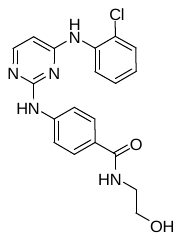  -logIC_50_ **7.394** | 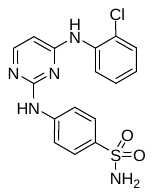  -logIC_50_ **7.364** | 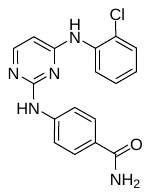  -logIC_50_ **7.299** | 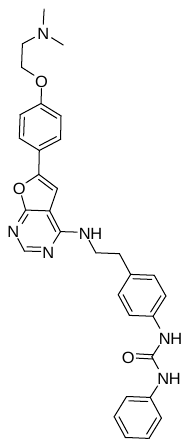  -logIC_50_ **7.276** | 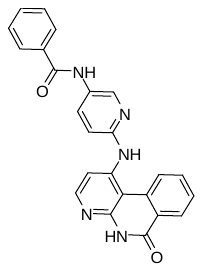  -logIC_50_ **7.102** |
| 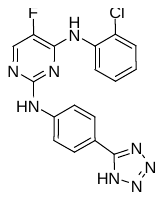  -logIC_50_ **7.077** | 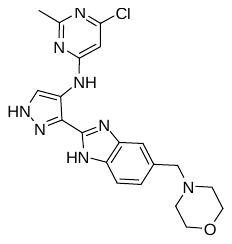  -logIC_50_ **7.036** | 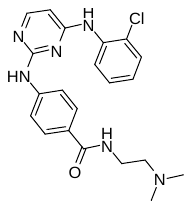  -logIC_50_ **7.027** | 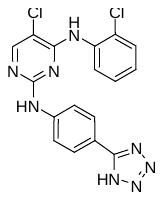  -logIC_50_ **6.975** | 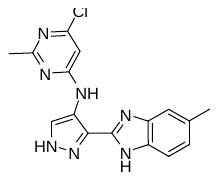  -logIC_50_ **6.857** |
| 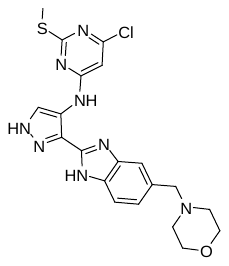  -logIC_50_ **6.842** | 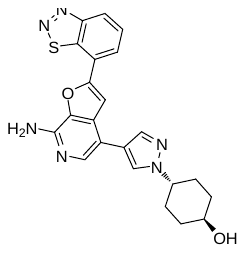  -logIC_50_ **6.796** | 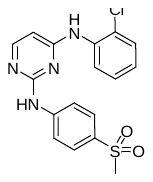  -logIC_50_ **6.745** | 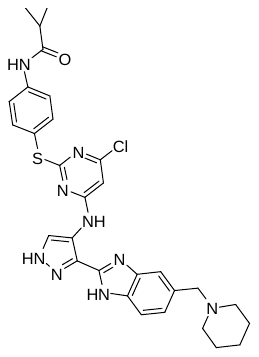 -logIC_50_ **6.719** | 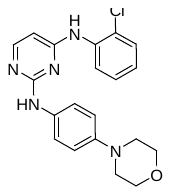  -logIC_50_ **6.712** |
| 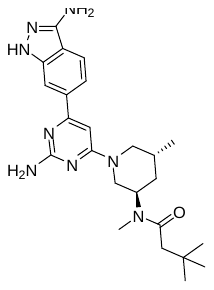  -logIC_50_ **6.699** | 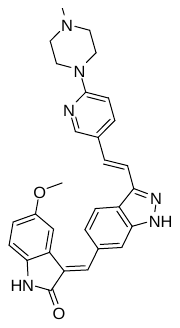  -logIC_50_ **6.678** | 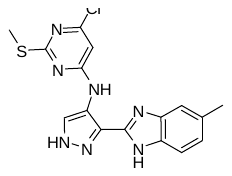  -logIC_50_ **6.636** | 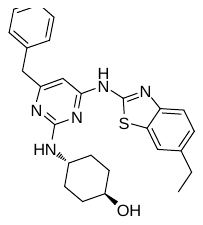  -logIC_50_ **6.400** | 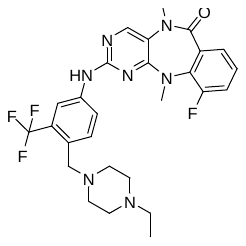  -logIC_50_ **6.312** |
| 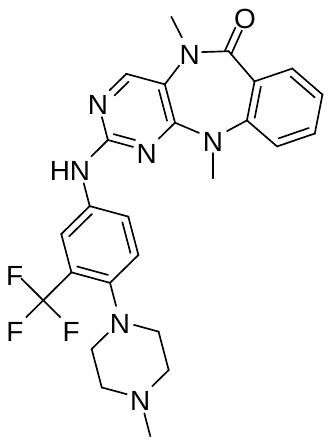  -logIC_50_ **6.111** | 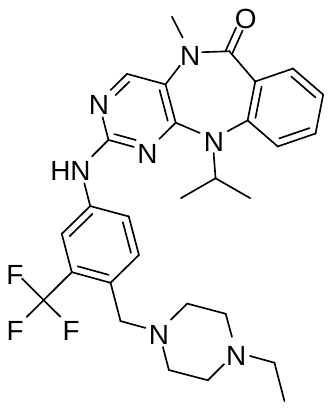  -logIC50 **6.107** | 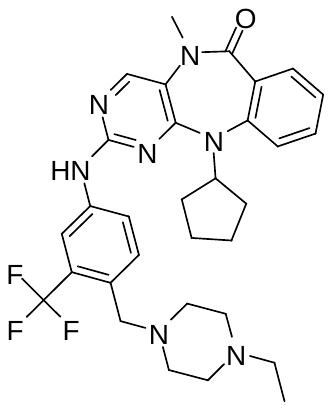  -logIC_50_ **6.013** | 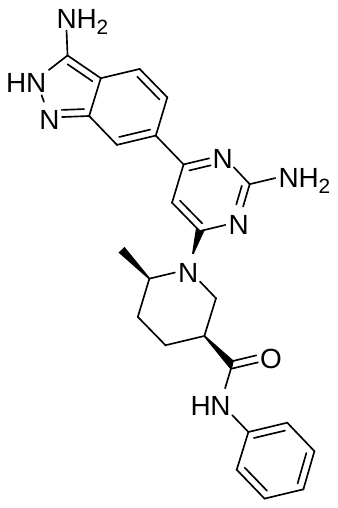  -logIC_50_ **6.000** | 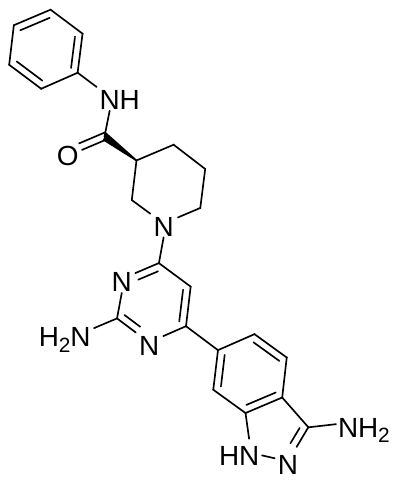 -logIC_50_ **5.900** |

**Table S2.** Chemical structures of test set compounds.

**(Source:**http://www.bindingdb.org/jsp/dbsearch/PrimarySearch_ki.jsp?tag=tg&kiunit=nM&icunit=nM&column=IC50&submit=Search&energyterm=kJ%2Fmole&target=Aurora+kinase+B)

| 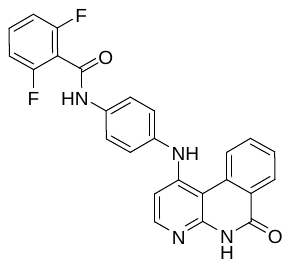  -logIC_50_ **9.222** | 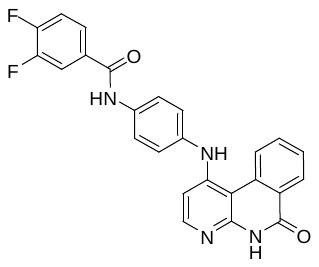  -logIC_50_ **9.097** | 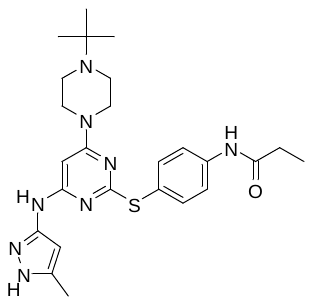 -logIC_50_ **7.959** | 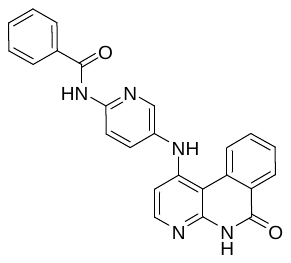  -logIC_50_ **7.886** |
| --- | --- | --- | --- |
| 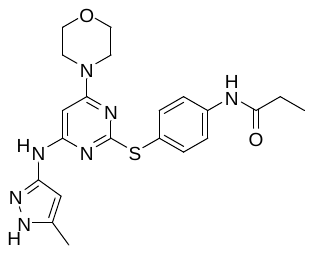  -logIC_50_ **7.854** | 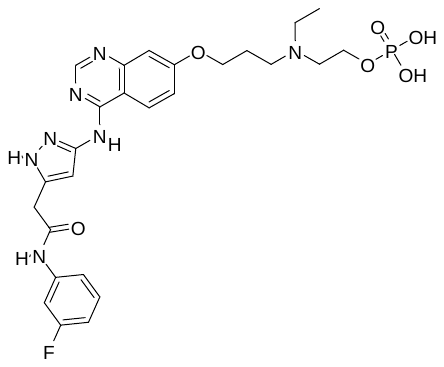  -logIC_50_ **7.810** | 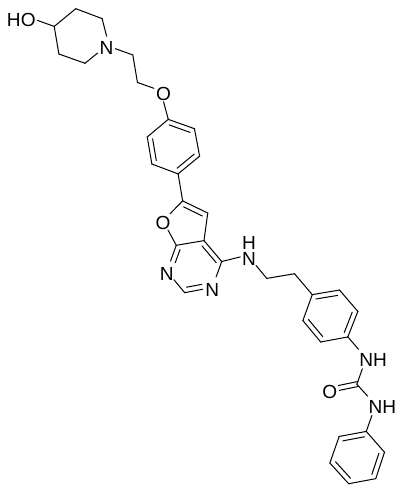  -logIC_50_ **7.770** | 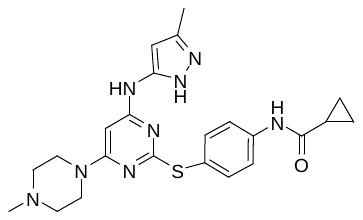  -logIC_50_ **7.703** |
| 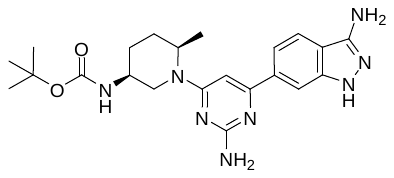  -logIC_50_ **7.300** | 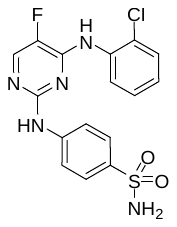  -logIC_50_ **7.247** | 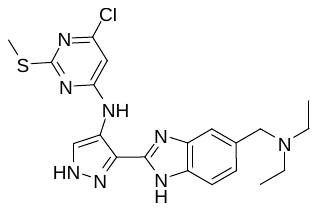 -logIC_50_ **7.004** | 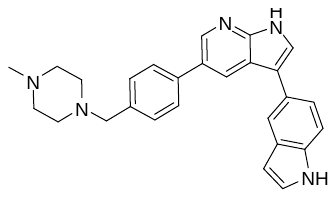  -logIC_50_ **6.910** |
| 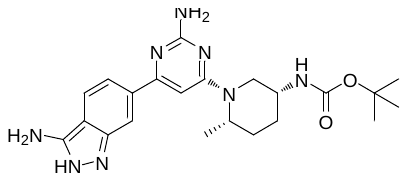  -logIC_50_ **6.800** | 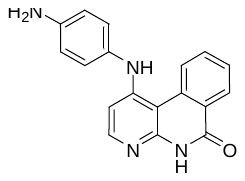  -logIC_50_ **6.759** | 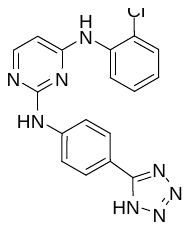  -logIC_50_ **6.646** | 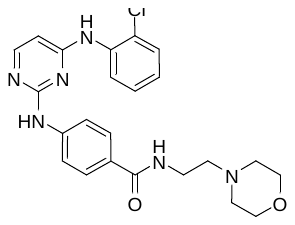  -logIC_50_ **6.635** |
| 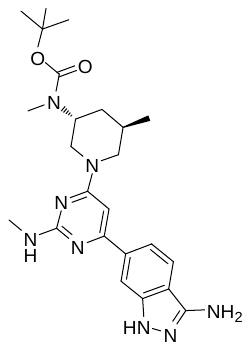  -logIC_50_ **6.600** | 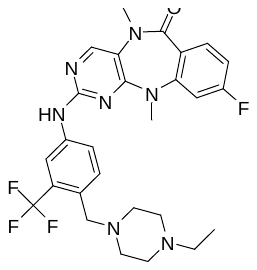  -logIC_50_ **6.232** |  |  |

**S1 Fig.** 2-D view of Interactions of (a) Compound 1, (b) Compound 2, (c) Compound 3, (d) Compound 4, (e) Compound 5 with 4AF3 protein.

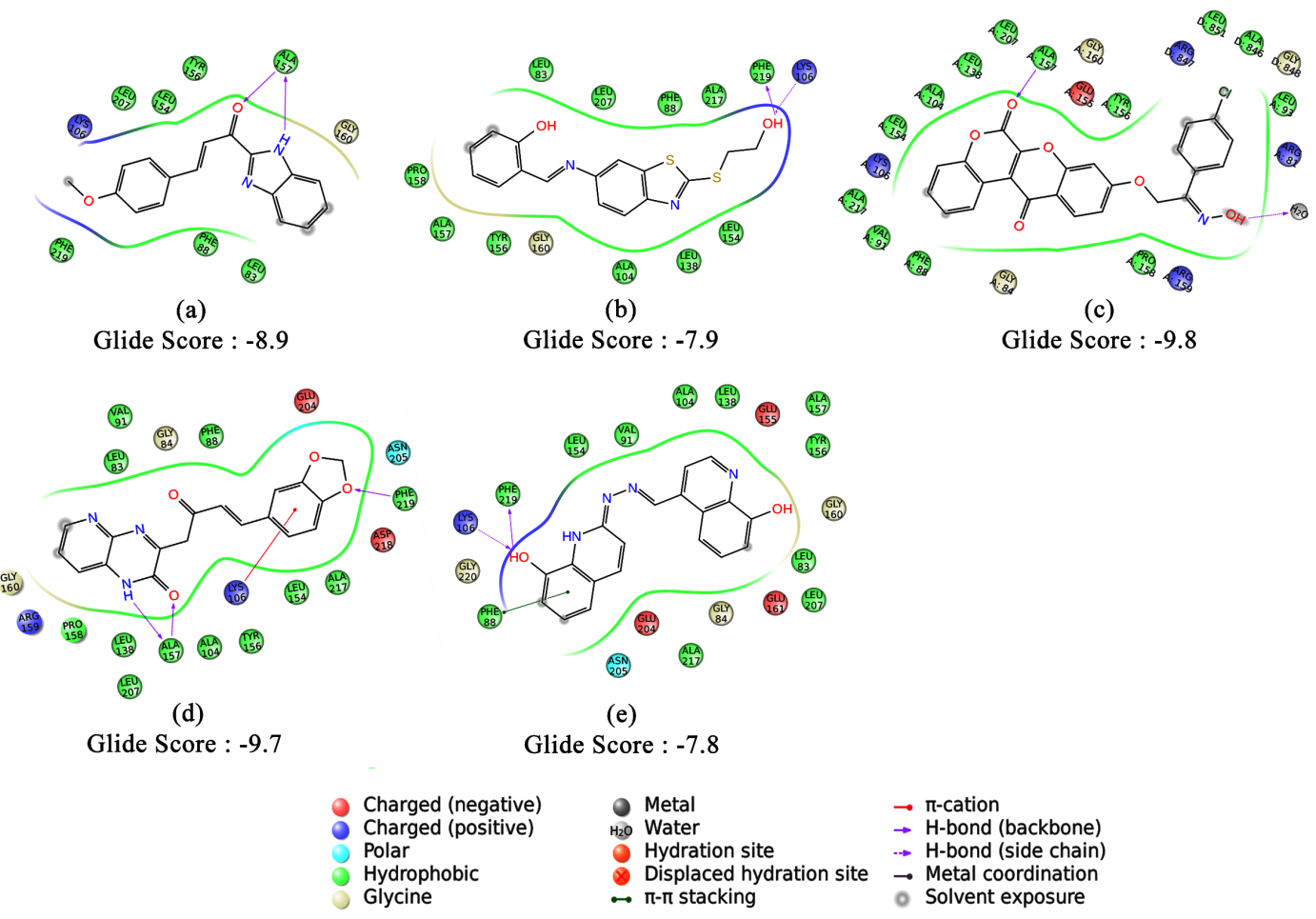

**S2 Fig.** Docking poses of Aurora−B kinase inhibitors (a) Hesperadin, (b) VX−680 and (c) ZM447439

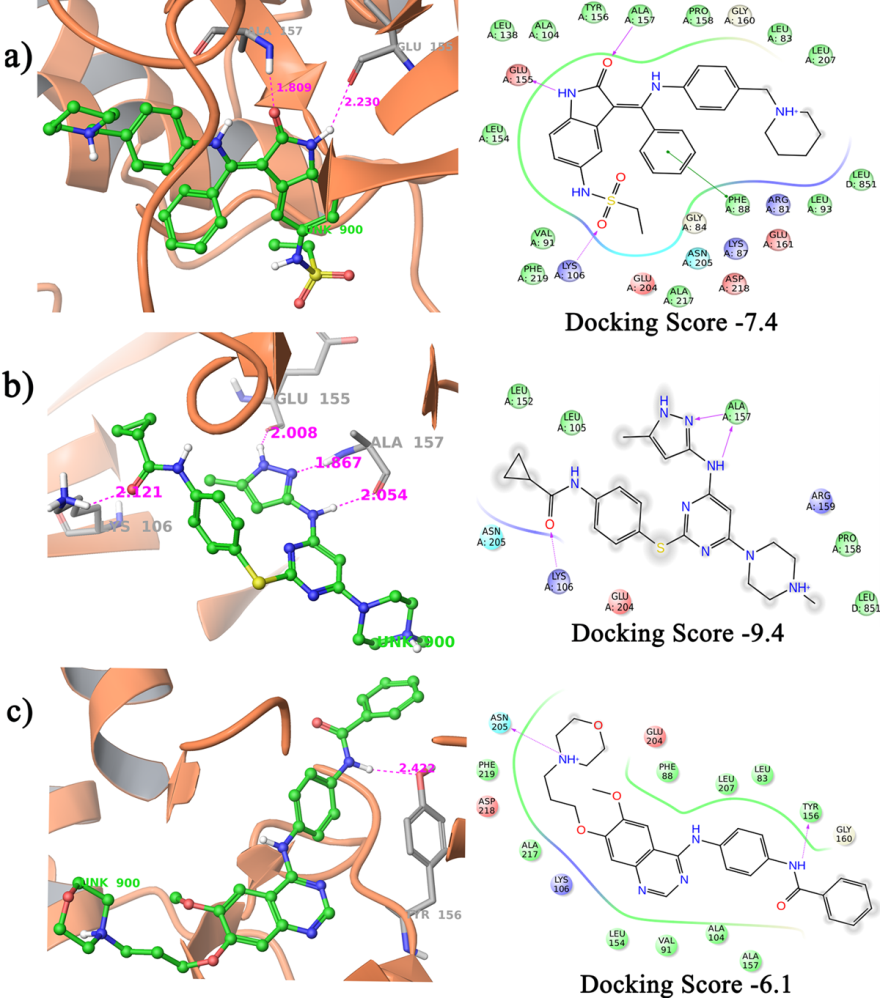
